# A chemoproteomic platform for reactive fragment profiling against the deubiquitinases

**DOI:** 10.1101/2023.02.01.526632

**Authors:** Rosa Cookson, Aini Vuorinen, Jonathan Pettinger, Cassandra R. Kennedy, Joanna M. Kirkpatrick, Rachel E. Peltier-Heap, Andrew Powell, Ambrosius P. Snijders, Mark Skehel, David House, Katrin Rittinger, Jacob T. Bush

**Affiliations:** Crick-GSK Biomedical LinkLabs, GSK, Gunnels Wood Road, Stevenage, Hertfordshire, SG1 2NY, UK; Molecular Structure of Cell Signalling Laboratory, The Francis Crick Institute, 1 Midland Road, London, NW1 1AT, UK; Proteomics Science Technology Platform, The Francis Crick Institute, 1 Midland Road, London, NW1 1AT, UK

**Keywords:** Deubiquitinases, reactive fragments, chemoproteomics, activity-based protein profiling, mass spectrometry, fragment-based drug discovery

## Abstract

Chemoproteomics is a powerful method capable of detecting interactions between small molecules and the proteome, however its use as a high-throughput screening method for chemical libraries has so far been limited. To address this need, we have further developed a chemoproteomics workflow to screen cysteine reactive covalent fragments in cell lysates against the deubiquitinating (DUB) enzymes using activity-based protein profiling. By using targeted ubiquitin probes, we have addressed sensitivity and affinity limitations, enabling target identification and covalent fragment library profiling in a 96-well plate format. The use of data independent acquisition (DIA) methods for MS analysis combined with automated Evosep liquid chromatography (LC) reduced instrument runtimes to 21 minutes per sample and simplified the workflow. In this proof-of-concept study, we have profiled 138 covalent fragments against 57 DUB proteins and validated four hit fragments against OTUD7B and UCHL3 through site identification experiments and orthogonal biochemical activity assays.

**Graphical abstract:** 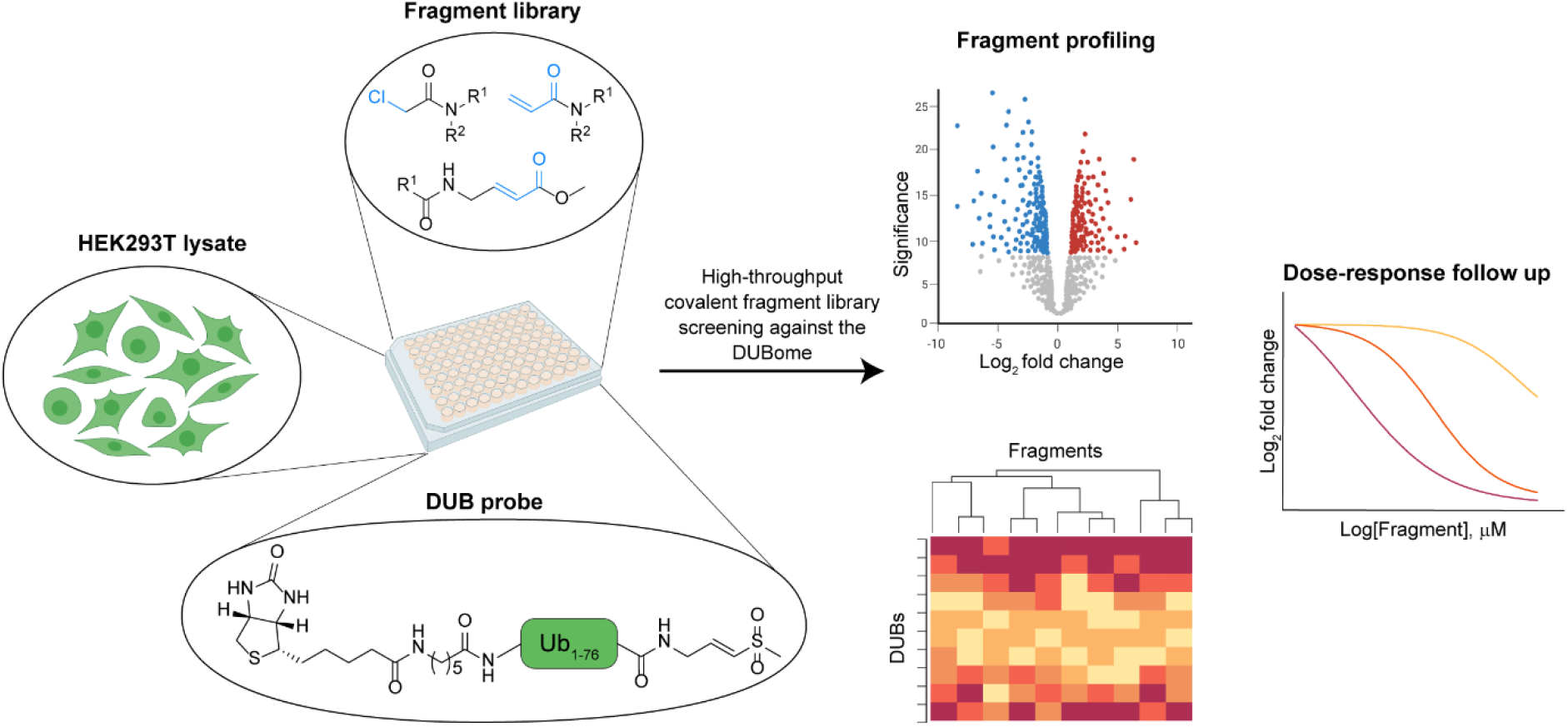

## Introduction

The development of chemical probes for the entire human proteome is an ambitious goal of the research community that will have profound impact on our understanding of protein function and human disease, ultimately translating into new innovative medicines.^1–3^ Currently, ligands exist for approximately 10% of the proteome, and just 22-40% is estimated to be druggable by traditional small molecules.^4, 5^ Recent advances in genomic approaches are highlighting novel, understudied targets from the human genome, and chemical probes provide an essential method to enable rapid validation and prioritisation of these targets.^6–11^

Traditional small molecule screening has so far been well utilised for chemical probe discovery, however, in order to accelerate this discovery, new orthogonal technologies are required.^12–16^ One such opportunity is the screening of reactive fragment libraries through chemoproteomic workflows.^17–19^ Fragments allow efficient coverage of chemical space with small libraries, while the functionalisation of the fragment with a reactive group affords increased potency and selectivity, permitting robust detection of fragment-protein interactions in cells (Figure 1A). Reactive fragment screening is typically performed with purified protein and has had significant impact in this field, ^19–26^ even enabling targeting of shallow protein pockets. Examples of fragment-derived medicines include inhibitors of KRas^G12C^ (Sotorasib),^27^ tumour target Pin1,^28^ and the SARS-CoV-2 target M^pro^.^29^

**Figure 1.**
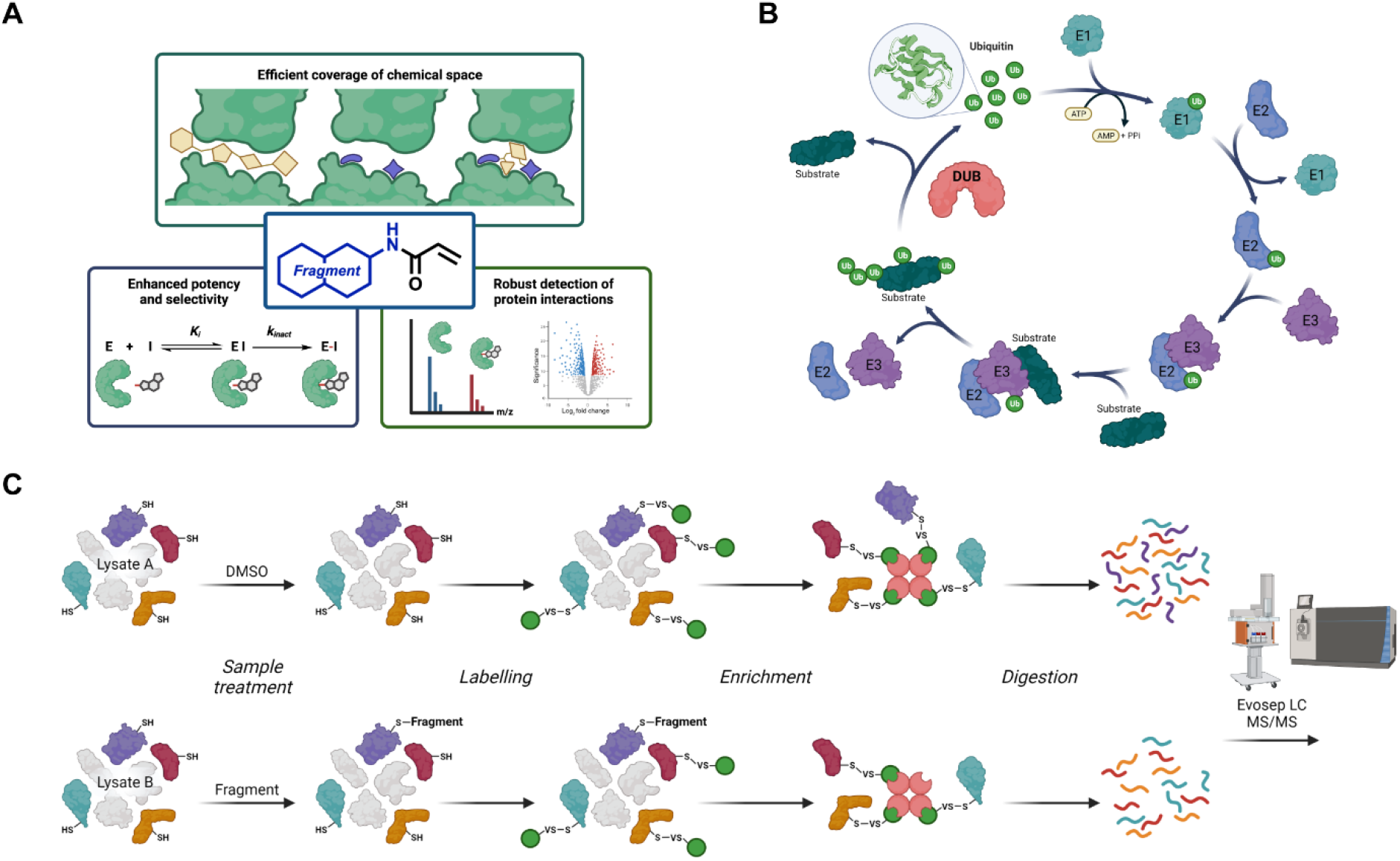
Fragment-based drug discovery of DUBs using chemoproteomics. A)Key advantages of screening with reactive fragments. B)The ubiquitin cycle, with ubiquitin (green), E1-activating enzymes (turquoise), E2-conjugating enzymes (blue), E3 ligases (purple), substrates (dark green) and deubiquitinase enzymes (pink). C)Schematic of the activity-based protein profiling workflow. Lysates are treated with covalent fragments or DMSO, followed by treatment with an activity-based DUB probe. Labelled proteins are enriched, digested and analysed by LC-MS/MS.

Forthcoming advances will likely take in fragment-based drug discovery from an *in vitro* screening method to an *in cellulo* platform capable of simultaneously detecting and studying fragment-protein interactions across the proteome.^17, 18, 30, 31^ Importantly, this would enable profiling of target engagement of endogenous proteins in their native context, where protein-protein interactions, post-translational modifications and subcellular localisation alter their activities and ligandability. Furthermore, such a platform allows the study of proteins that are difficult to isolate recombinantly for *in vitro* screening and assays.

So far, a key limitation to developing this reactive fragment screening platform in cells has been the sensitivity of mass spectrometry (MS) analysis, which makes achieving sufficient coverage of functionally relevant protein sites a challenge. Despite rapid advances in profiling cysteine reactive fragments against the ‘cysteinome’ using iodoacetamide based probes,^32, 33^ only ~10% of the cysteinome can currently be liganded with fragments. One approach to circumvent this is to use targeted probes against a protein family of interest, for example activity-based probes (ABPs) which target a conserved pocket or protein surface. Employing these ABPs as reporter molecules in a chemoproteomics increases the depth of protein coverage within the desired protein family when compared to a cysteinome-wide approach to a level where fragment screening can be employed effectively.

A second barrier for chemoproteomic screening of chemical libraries is the time required for liquid chromatography (LC), which can typically be up to three hours per sample. Significant improvements have been achieved through isotope labelling, enabling multiplexing and analysis of 10-15 samples in one run.^32, 34, 35^ However, isotope labelling reagents are expensive and therefore not ideally suited to screening large numbers of compounds. In addition, data-dependent acquisition (DDA) is the most frequently utilised MS method in chemoproteomics, but this can limit the comparability between MS runs, a necessity for robust profiling of a library. Recent advances in data independent acquisition (DIA) and label free quantification methods have enabled shorter LC run times, and have reduced the number of missing values, relative to DDA,^36–38^ offering great potential for library profiling.^36, 37, 39^

Herein we describe a high throughput chemoproteomics platform to screen cysteine reactive fragment libraries against deubiquitinases (DUBs). Ubiquitination is a critical and complex post-translational modification which controls and regulates most cellular processes. The ubiquitin cycle is carefully maintained by ubiquitin system proteins: substrate ubiquitination is a result of an enzymatic cascade involving E1-activating enzymes, E2-conjugating enzymes, and E3 ligases; substrate deubiquitination is controlled by DUBs (Figure 1B).^23^ In humans, DUBs can be classified into two categories: the metalloproteases (JAMM family) and cysteine proteases (USP, OUT, MJD, UCH, MINDY and ZUFSP families). Both categories of DUBs have been implicated in many diseases including central nervous system (CNS) disorders, inflammation, immunity and infectious diseases,^40^ and have been ardently studied to decipher their mechanisms and substrate specificities. Despite this, new chemical probes are still needed to explore the biology of DUB proteins and aid development of therapeutics.^41^

In this study we describe the development of a workflow that was optimised in 96-well plate format and employs DIA-MS analysis to enable fast analysis times while profiling >50% of the native DUBome (Figure 1C). This platform was used to screen a library of 138 fragments, identifying functionally relevant hits for numerous members of the DUB family and demonstrating the utility of the approach in expanding the liganded proteome.

## Results

### Activity-based probe selection and chemoproteomic platform development

To establish a platform for screening cysteine-reactive fragments against the endogenous DUBome, a DUB specific ABP was first selected. DUB-targeting ABPs mimic native ubiquitin but are modified at the ubiquitin C-terminus with an electrophilic warhead. When bound to cysteine protease DUBs, this warhead irreversibly labels the active site cysteines.^42, 43^ An affinity handle, such as biotin, at the N-terminus of ubiquitin enables downstream enrichment of labelled proteins. We selected three DUB ABPs with different electrophiles to identify which would give the best coverage and enrichment of the DUBome: biotinylated ubiquitin-vinylsulfone (Biotin-Ahx-Ub-VS), biotinylated ubiquitin-propargylamide (Biotin-Ahx-Ub-PA) and biotinylated ubiquitin-vinyl methyl ester (Biotin-Ahx-Ub-VME) (Figure 2A).

**Figure 2.**
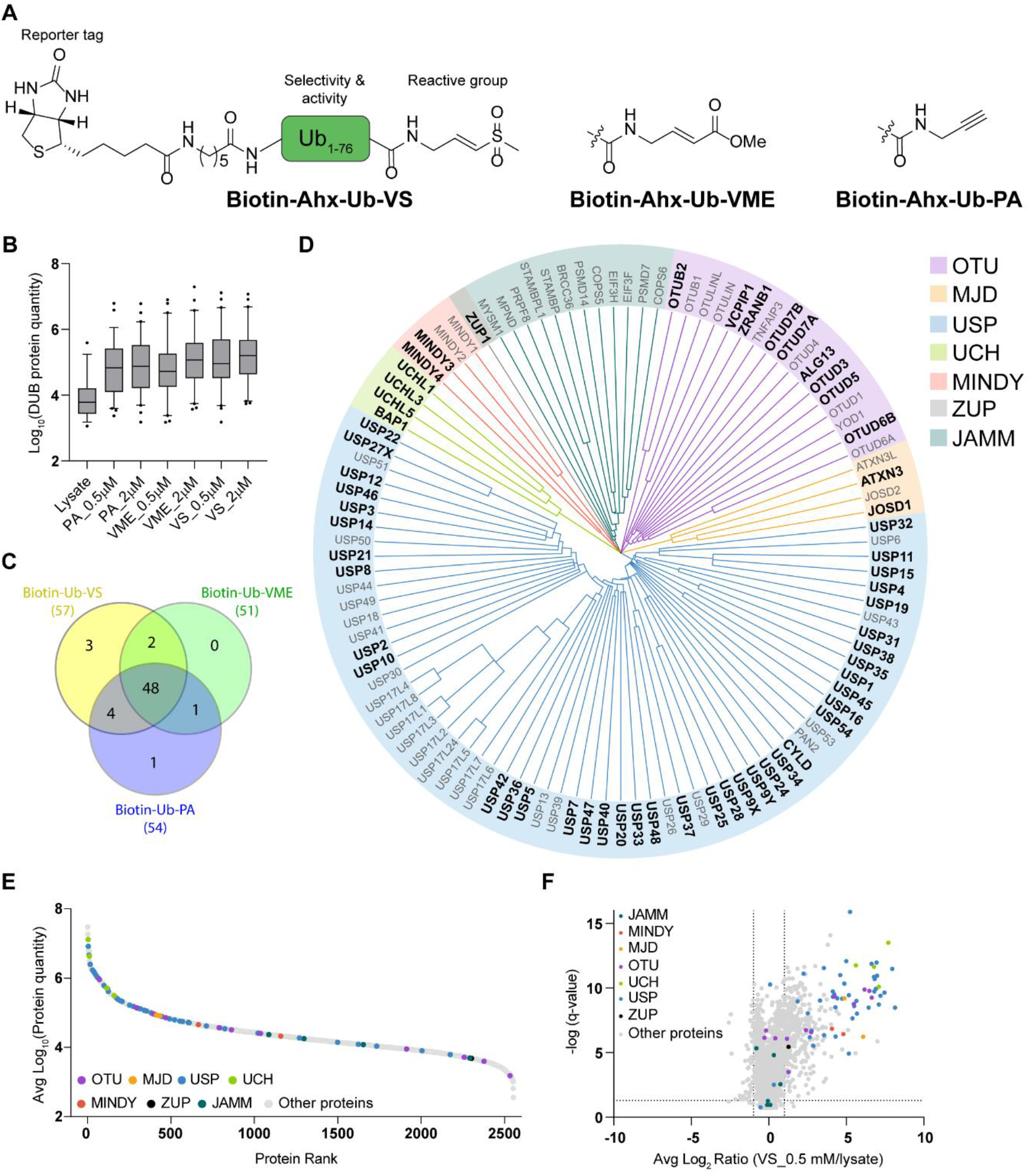
Method development and DUBome coverage using automated Evosep One LC and Orbitrap Fusion Lumos MS alongside DIA for MS analysis. A) Structures of activity-based DUB probes Biotin-Ahx-Ub-VS, Biotin-Ahx-Ub-VME and Biotin-Ahx-Ub-PA. B) DUB protein quantities identified by each activity-based DUB probe (VS = Biotin-Ahx-Ub-VS; VME = Biotin-Ahx-Ub-VME; PA = Biotin-Ahx-Ub-PA). Box plot shows the median central line and extends from 25th to 75th percentiles. Whiskers represent protein quantity within 5th to 95th percentiles. C) Venn diagram showing overlap of significantly enriched DUBs for Biotin-Ahx-Ub-VS, Biotin-Ahx-Ub-VME and Biotin-Ahx-Ub-PA probes at 0.5 μM when compared to untreated lysate. D) DUB phylogenetic tree highlighting in bold the significantly enriched DUBs when Biotin-Ahx-Ub-VS (0.5 μM) treatment is compared to untreated lysate (avg log_2_ ratio ≤ −1, q-value ≤ 0.05). Full length DUB sequences were aligned with COBALT^44^ and subsequently visualised with iTOL.^45^ E) Protein rank plot for Biotin-Ahx-Ub-VS treated sample (0.5 μM) listing an average protein quantity for all quantified proteins in descending values. DUBs are coloured based on their subfamilies. F) Volcano plot showing significantly enriched proteins (avg log_2_ ratio ≤ −1, q-value ≤ 0.05) when Biotin-Ahx-Ub-VS (0.5 μM) treated sample is compared to untreated lysate. DUBs are coloured based on subfamilies.

We first tested the DUB ABPs in a proteomics workflow that was designed to be fully applicable for subsequent downstream fragment library screening. To reduce running costs and negate the need for multiplexing, simplifying the workflow, we chose to use DIA MS analysis. Furthermore, we aimed to reduce the MS instrument runtime as much as possible. HEK293T lysates were treated with each of the three probes at two concentrations (0.5 and 2 μM) and DMSO for one hour, before labelled proteins were enriched using avidin and digested with Trypsin. Peptides were analysed by LC-MS/MS using an Evosep One – Orbitrap Fusion Lumos with a 44 minute LC gradient and a DIA method.^39^

Each probe afforded similar protein intensities, coefficients of variation (CV) and high sequence coverage of the DUBs (Figure 2B and SI Figure 1B and E). All three probes were found to enrich 51-57 DUBs compared to the DMSO control (SI Figure 1A), representing 55-61% of the catalytic cysteine-containing DUB family. Of these, 48 were enriched across the three probes, and Biotin-Ahx-Ub-VS showed the best coverage (Figure 2C) and was selected for future experiments. Furthermore, Biotin-Ahx-Ub-VS showed coverage across each DUB subfamily (Figure 2D), with the exception of the metalloprotease JAMM DUB family which are out of scope for these probes. In addition to selective enrichment compared to the DMSO control, DUB proteins were ranked highly in absolute intensity, highlighting the robust levels of enrichment over the proteome background (Figure 2E). Biotin-Ahx-Ub-VS exhibited high selectivity for the identified cysteine protease DUBs (median avg log_2_ ratio = 4.86) over the 2502 non-DUB enriched proteins (median avg log_2_ ratio = −0.04) (Figure 2F). A lower concentration of Biotin-Ub-VS probe at 0.2 μM was explored, however, while the CVs remained similar, this led to a decrease in DUB protein intensities (SI Figure 1C and D).

We subsequently sought to increase the throughput of the analysis by decreasing the LC runtimes. Three LC gradients were compared, via 44-, 21- and 11-minute methods. Minimal reduction in the intensities and CVs of DUBs was observed upon reduction of run-time, suggesting all methods would be suitable for library screening (SI Figure 2A, B).

### Validation of chemoproteomics workflow with known inhibitors

To validate our protocol in a displacement assay, we selected three reported DUB ligands for initial profiling; the parent of Biotin-Ahx-Ub-VS, Ub-VS,^46^ the pan-DUB inhibitor PR619,^47^ and a Ubiquitin Specific Protease (USP) inhibitor (herein referred to as ‘USP probe’) that targets 12 USPs.^48^ HEK293T lysates were incubated with inhibitor (2.5 μM Ub-VS, 50 μM PR619, or 50 μM USP probe) or DMSO vehicle for three hours prior to treatment with Biotin-Ub-VS probe (0.5 μM) and subsequent enrichment and MS-analysis as described above. This was performed with six replicates to thoroughly determine reproducibility.

Ub-VS outcompeted 42 out of 53 DUBs enriched by the Biotin-Ahx-Ub-VS ABP (q-value ≤ 0.05 and avg log_2_ ratio ≤ −1, Figure 3A), indictive of mechanism-based capture. PR619 and USP probe outcompeted 27 and 35 DUBs respectively (q-value ≤ 0.05 and average log_2_ ratio ≤ −1, Figures 3B and 3C). Of these, 21 DUBs were competed by both compounds. The reproducibility of the platform was found to be excellent, with 49 out of 51 DUBs quantified in all six replicates (DMSO control) and median CVs for the DUB proteins were between 24-29% (SI Figures 3A and B). As a result of the high reproducibility, the number of biological replicates was reduced to three in subsequent experiments.

**Figure 3.**
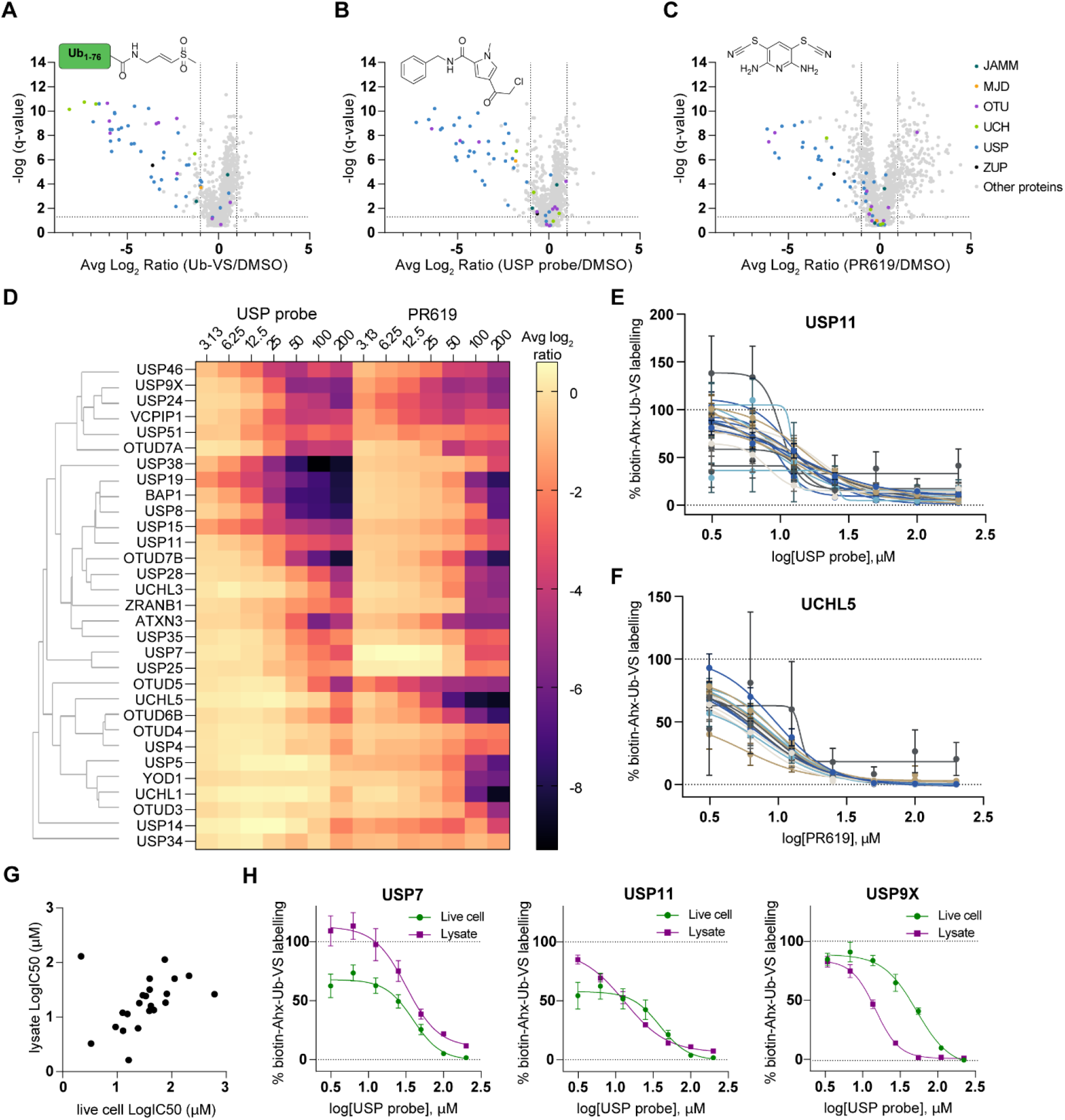
Validation of the chemoproteomics workflow with known DUB inhibitors. A) Volcano plot showing significantly competed proteins (avg log_2_ ratio ≤ −1, q-value ≤ 0.05) when Ub-VS (2.5 μM) treated sample is compared to DMSO (Biotin-Ahx-Ub-VS, 0.5 μM). DUBs are coloured based on subfamilies. B) Volcano plot showing significantly competed proteins (avg log_2_ ratio ≤ −1, q-value ≤ 0.05) when USP probe (50 μM) treated lysate is compared to DMSO (Biotin-Ahx-Ub-VS, 0.5 μM). DUBs are coloured based on subfamilies. C) Volcano plot showing significantly competed proteins (avg log_2_ ratio ≤ −1, q-value ≤ 0.05) when PR619 (50 μM) treated lysate is compared to DMSO (Biotin-Ahx-Ub-VS, 0.5 μM). DUBs are coloured based on subfamilies. D) Heatmap showing dose-response chemoproteomics data for a panel of DUBs (lysate treatment). Avg log_2_ ratios of inhibitor/DMSO are plotted for each DUB. DUBs are clustered on the y-axis by single linkage clustering method using cosine correlation distance measure. E) Dose-response chemoproteomics data for USP11 peptides. Data is presented as mean ± SEM, *n* = 3. The curves were fitted with GraphPad Prism 9 using four parameter nonlinear regression. F) Dose-response chemoproteomics data for UCHL5 peptides. Data is presented as mean ± SEM, *n* = 3. The curves were fitted with GraphPad Prism 9 using four parameter nonlinear regression. G) Scatter plot showing logIC50 values for a panel of DUBs comparing USP probe treated lysate and live cell dose-response chemoproteomics data. logIC50 values were extracted from dose-response curves that were fitted with GraphPad Prism 9 using four parameter nonlinear regression. H) Selected examples of dose-response chemoproteomics data to compare lysate and live cells treated with USP probe. Data is presented as mean ± SEM, *n* = 3. The curves were fitted with GraphPad Prism 9 using four parameter nonlinear regression. All dose-response curves are presented in SI Figure 4.

To further validate the workflow for screening compound libraries against the DUBome, the two DUB inhibitors, PR619 and USP probe, were profiled in a seven-point dose-response experiment, with concentration responses observed at both the protein and peptide level (Figure 3D, E and F). Clear differences were observed for the DUB interaction profiles of the two compounds. A greater number of IC_50_ values were determined for the USP probe (32) compared to PR619 (26), consistent with the initial single concentration experiment (SI Table 1). As expected, the pan-DUB inhibitor PR619 exhibited activity across a broad spectrum of DUBs: IC_50_ values below 10 μM were observed for OTUD5, USP46, VCPIP1, USP9X and UCHL5. The USP inhibitor showed increased selectivity for the USP proteins at lower concentrations, with IC_50_ values below 10 μM for USP19, USP15, USP38, USP46, BAP1 and USP8.^48^

### Comparison of inhibitor treatments in lysates and in live cells

Following workflow optimisations in cell lysates, we profiled the USP probe in a live cell dose response experiment. HEK293T cells were incubated for three hours with USP probe (0 - 200 μM), and cells were lysed prior to addition of Biotin-Ahx-Ub-VS probe (0.5 μM). Labelled proteins were enriched and MS-analysis was performed as described previously. Dose response curves were obtained for 37 DUBs, consistent with the lysate-based protocol. However, there were some notable differences in which DUBs were detected. We speculate that this is a consequence of the respective stability of these DUBs in the two different workflows. Importantly, there was a good correlation between IC_50_ values observed from the cell and lysate treatments, providing confidence that lysate-based screening is a reasonable proxy for engagement of proteins in cells, while enabling higher throughput (Figure 3G, H and SI Figure 4). Nevertheless, cell-based experiments would always be recommended for validation of any interactions identified in lysate-based screening.

### Cysteine reactive fragment library screening against the DUBome

Following the establishment and validation of this new and powerful high throughput chemoproteomics platform for the DUBome, we next wanted to perform fragment screening with our methodology. A cysteine reactive fragment library was screened to identify new starting points for the development of covalent DUB inhibitors (Figure 4A). A library of 138 cysteine reactive fragments was selected, comprised of three electrophilic groups (chloroacetamide, acrylamide and methyl acrylate) linked to a chemically diverse set of fragments to assess a range of reactivities, geometries, and functionality (Figure 4A, SI Figure 5A). The library of fragments had molecular weights of between 180 and 400 g/mol (SI figure 5B). The fragments were incubated in cell lysates at 200 μM for three hours and enriched DUBs were analysed using a 21 minute LC method. This enabled a throughput of 60 samples per day, with the entire screen requiring just 11 days of instrument time. These 11 days included a number of quality controls (QCs) and blanks to ensure the quality of the data. Excellent consistency in the performance of the protocol and instrument performance was observed across all plates and the multiple days of data acquisition (SI Figure 6).

**Figure 4.**
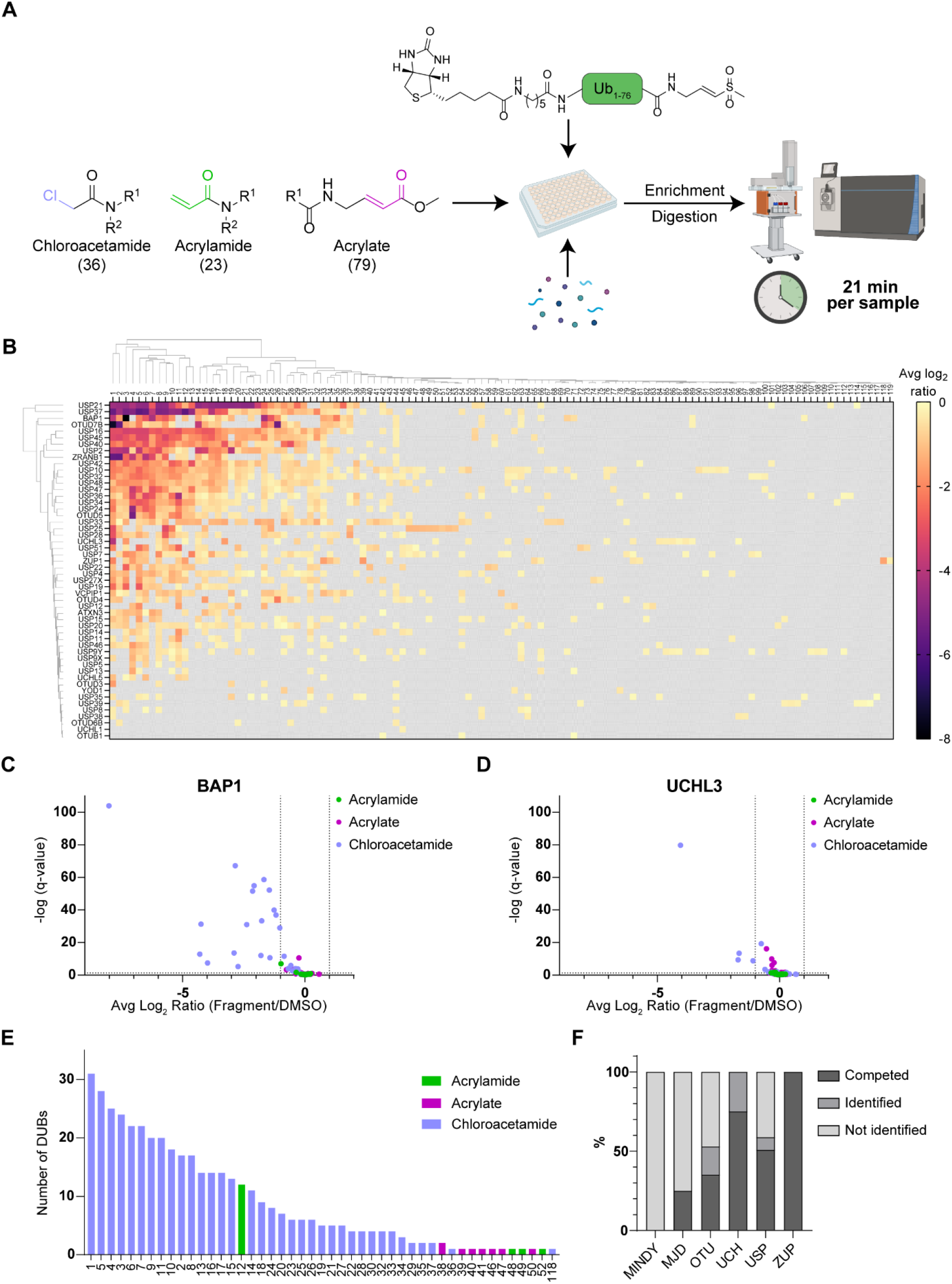
Fragment library screening. A) Schematic overview of reactive fragment library screening by chemoproteomics. Lysates were treated with the reactive fragments, followed by Ub-VS incubation, enrichment of labelled proteins and digestion for LC-MS/MS analysis. B) Heatmap showing fragment library screening results. Avg log_2_ ratio of fragment/DMSO ≤ 0 is plotted for each DUB when q-value ≤ 0.05. Fragments and DUBs are clustered by UPGMA clustering method using Euclidean distance measure. C) Volcano plot showing all fragments avg log_2_ ratio and q-value for all fragments (200 μM) against BAP1. D) Volcano plot showing all fragments avg log_2_ ratio and q-value for all fragments (200 μM) against UCHL3. E) Number of DUBs competed by each fragment when avg log_2_ ratio ≤ −1 and q-value ≤ 0.05. Bars are coloured based on electrophile. F) Percentage of DUBs in each family that were competed (avg log_2_ ratio ≤ −1, q-value ≤ 0.05), identified and not identified.

The resulting full-matrix data set gave competition ratios to describe the interaction between 119 fragments and 52 endogenous DUBs (average log_2_ ratio ≤ 0, q-value ≤ 0.05 and peptides ≥ 2) (Figure 4B). The chloroacetamides were found to exhibit increased promiscuity (as seen by clustering to the lefthand side of the heatmap) in line with their reactivity, while the acrylates and acrylamides were found to interact with fewer DUBs (as seen by clustering to the righthand side of the heatmap). Fragments interactions were filtered for with average log_2_ ≤ −1 (> 50% competition). Some proteins were found to interact with multiple fragments, such as BAP1 and USP21 (Figure 4C and SI Figure 7), whereas other DUBs interacted with few fragments, such as UCHL3 and USP24 (Figure 4D and SI Figure 7). In total, 43 DUBs were competed by 47 fragments (Figure 4E). At least one DUB interaction was found for 35 of the 36 chloroacetamides (97%) screened. This is in comparison to 17% of acrylamides and 10% of acrylates. *p*-Nitrophenyl acrylamide 12 sits as an outlier of the acrylamides as it significantly competes 12 DUB proteins, which is consistent with the enhanced reactivity induced by the electron deficient phenyl ring. There was good coverage across all DUB sub-families, with covalent fragments found to compete USP, MJD and OTU families (59, 25 and 53% respectively, Figure 4F). The highest proportion of covalent hits were observed for the smaller sub-families UCH and ZUP, with covalent fragments found to compete 75% and 100% members of the families, respectively.

### Validation of fragment-DUB interactions

Four hit fragments were selected for dose-response validation based on high significance scores, competition ratios and a range of selectivity profiles (Figure 5A, B and SI Figure 8A, B). Fragments 1, 2 and 3 labelled multiple DUB proteins, and clustered together to the lefthand side of the heat map. Fragment 26 was selected for further analysis based on its selective labelling profile, even at 200 μM. Two of the DUBs significantly competed by fragment 26 (BAP1 and OTUD7B) were also significantly competed by fragments 1-3.

**Figure 5.**
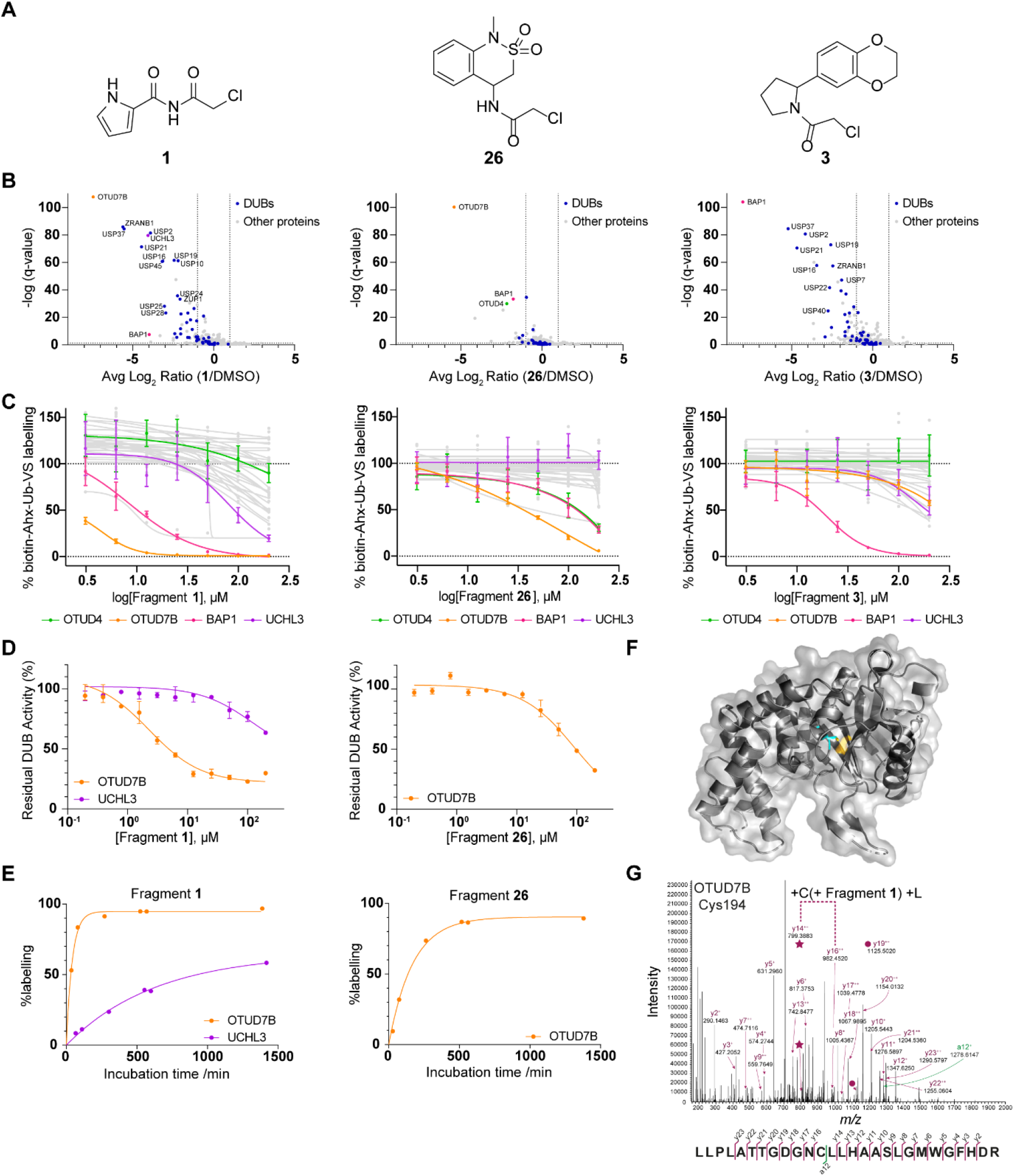
Hit fragment validation. A)Structures of selected hit fragments **1**, **3** and **26**. B)Volcano plots showing significantly competed proteins (avg log_2_ ratio ≤ −1, q-value ≤ 0.05) when hit fragment (200 μM) treated samples are compared to DMSO (Biotin-Ahx-Ub-VS, 0.5 μM). DUBs are highlighted in blue. C)Dose-response chemoproteomics data for DUB proteins. Data is presented as mean ± SEM, *n* = 3. The curves were fitted with GraphPad Prism 9 using four parameter nonlinear regression. D)Enzymatic inhibition assay data for hit fragments **1** and **26**. Data is presented as mean ± SD, *n* = 3. The curves were fitted with GraphPad Prism 9 using three parameter nonlinear regression. E)Intact-protein LCMS time-course percentage labelling data for fragments **1** and **26**. The curves were fitted with GraphPad Prism 9 using one phase association. F)Structure of OTUD7B highlighting the active site Cys194 (PDB: 5LRU).^49^ G)LC-MS/MS spectra of the peptide _183_LLPLATTGDGNCLLHAASLGMWGFHDR_209_ crosslinked to fragment **1** indicating OTUD7B Cys194 as the fragment binding site.

The four fragments were further profiled in the chemoproteomics DUB screening platform at a range of concentrations from 3.1 - 200 μM (Figure 5C and SI Figure 8). For each of these four fragments, the most significantly competed DUB protein in the single concentration experiment corresponded with the highest potency in the dose response. For fragments 1, 2 and 26 this was OTUD7B, and for fragment 3 it was BAP1. Each of the fragments showed engagement of BAP1, confirming that BAP1 and OTUD7B are both promiscuous proteins. Fragment 3 showed a high selectivity window for BAP1, which was not predicted from the single concentration experiment. Three DUBs significantly competed by fragment 26 (OTUD7B, BAP1 and OTUD4) in the original single concentration experiment were engaged in the dose-response with the same rank order selectivity (Figure 5B, C).

### Orthogonal assays for DUB target validation

To gain further confidence in the fragment-target pairings, the fragments were profiled in orthogonal assays. To assess inhibition of DUB activity by covalent modification, an activity-based enzymatic assay that follows cleavage of ubiquitin-rhodamine by recombinant protein was employed. Recombinant OTUD7B was incubated with fragments 1 and 26 (Figure 5D) and fragment 2 (SI Figure 8D) for three hours prior to the addition of Ub-rhodamine. Fluorescence intensities were plotted as a percent activity compared to DMSO controls. Consistent with chemoproteomics dose experiments, each fragment showed a dose dependent response, with the most potent interaction seen for fragment 1. The assay was repeated with recombinant UCHL3 and fragment 1 (Figure 5D), which was less potent in agreement with the chemoproteomic experiment. Unfortunately, BAP1 could not be assessed in this assay due to instability of the recombinant protein. To investigate kinetics of protein labelling with fragments 1, 2 and 26 we carried out a time-course of fragment engagement with recombinant OTUD7B and UCHL3 over 24 hours followed by LCMS analysis (Figure 5E and SI Figure 8E). The time dependent labelling of OTUD7B and UCHL3 correlated well with the enzymatic assays and chemoproteomic experiments. Labelling of UCHL3 by fragment 1 was significantly slower than of OTUD7B, explaining the less potent inhibition in the enzymatic assay.

Given the mechanism-based capture of the DUB activity-based probe it was expected that covalent modification of the fragment was occurring at the active site cysteine. In order to confirm this, site ID MS experiments were performed with UCHL3, OTUD7B and fragments 1, 2 and 26 (Figure 5F, G and SI Figure 8F, G and H). For all fragments, the predominant site of labelling was identified to be the catalytic cysteine of OTUD7B (Cys 194) across all fragments and UCHL3 (Cys 95) with fragment 1.

## Discussion

The development and discovery of chemical probes for the entire human proteome is recognised as a key objective that will accelerate our understanding of biology and disease. Ligand discovery traditionally relies on biochemical screening with purified protein, which can be time and cost intensive, and often does not effectively reflect *in vivo* ligandability. Covalent fragment-based ligand discovery *in cellulo* provides a powerful alternative due to the small size of libraries required and convenient readout of target engagement by LC-MS. However, to efficiently employ covalent fragment screening of entire libraries in a cellular environment requires the development of high throughput chemoproteomic workflows that are reliable and cost effective. Here we have reported the development of a powerful and efficient workflow and have used it to screen a library of 138 cysteine-reactive fragments against DUBs in an endogenous setting, enabling the identification of numerous fragment-protein interactions. Efficient chemoproteomic profiling of reactive fragment against the DUBs was achieved through optimisation of multiple steps. Typically, chemoproteomic workflows employ a purification or protein precipitation step, which complicates the workflow and limits parallel sample preparation. Through the use of a high affinity probe that was pre-conjugated to biotin, we removed the requirement for protein precipitation and established the protocol in 96-well plate format. The throughput of mass spectrometry analysis was optimised through the use of an Evosep LC system combined with label-free DIA methods to achieve instrument run times of 21 minutes per sample and a throughput of 60 samples per day, which we envisage could be further reduced to 11 minutes per sample. Advantages of this label free approach include facilitating comparison across hundreds of samples and reduction of costs associated with isotope labelling reagents. The workflow described here enabled reliable quantification of 57 DUBs with high reproducibility and low coefficients of variation.

Validation of the workflow as a screening platform was performed using three established DUB inhibitors, Ub-VS,^46^ PR619, ^47^ and a Ubiquitin Specific Protease (USP) inhibitor, USP probe. Interactions with multiple endogenous DUBs were reliably detected and concentration response experiments enabled differentiation of the potencies of each inhibitor-DUB interaction. Furthermore, screening of fragments in live cells afforded IC_50_ values that correlated well with those measured in lysates, supporting our approach of lysate screening as a suitable proxy for cells to improve the throughput of compound treatment.

Finally, we employed the platform to profile a reactive fragment library to identify ligands for proteins of the DUB family. A library of 138 reactive fragments was profiled against 52 endogenous DUBs, using just 11 days of instrument time. Hits were identified for 43 of the DUBs (>50% competition at 200 μM), and these may serve as starting points for the optimisation of more potent and selective inhibitors. Four hits were selected for profiling at a range of concentrations, with many interactions found to drop off quickly at lower concentration, but revealing relatively selective interactions for OTUB7B (IC_50_ < 3 μM) and BAP1 (IC_50_ ~10 μM). The potency and selectivity of these interactions correlated with covalent modification of purified proteins, as determined by intact mass spectrometry. Finally, inhibition of DUB activity was measured for OTUD7B and UCHL3, which was found to match the profile measured by chemoproteomics.

This chemoproteomics platform demonstrates a cost and time effective method for identifying fragments which can be further developed into potent and specific chemical probes against the DUBome. We envisage this platform could be used alongside AI/ML-based approaches for rapid compound optimisation. Furthermore, this adaptable workflow provides opportunities for screening additional protein classes and could be coupled with alternative reactive fragment warheads to target alternative amino acids.

## Acknowledgments

This work was supported by the Francis Crick Institute which receives its core funding from Cancer Research United Kingdom (CC 2075), the United Kingdom Medical Research Council (CC 2075), and the Wellcome Trust (CC 2075), by the Biotechnology and Biological Research Council, BB/T014547/1 to KR and DH, and by the Engineering and Physical Sciences Research Council, EP/V038028/1, to DH, JB and KR. For the purpose of Open Access, the author has applied a CC BY public copyright licence to any Author Accepted Manuscript version arising from this submission. Figures were created in BioRender.com under the institutional license belonging to the Francis Crick Institute.

## Author Contributions

K.R., D.H., A.P.S., and J.T.B. conceived the study. R.C., A.V., J.P., C.R.K., J.M.K., and R.E.P-H. designed and performed experiments. M.S. and A.P. supported proteomics and experimental work. R.C., A.V., C.R.K., J.P., K.R., and J.T.B. wrote the manuscript, and all authors approved the final version.

