## Supplementary Information for "A chemoproteomic platform for reactive fragment profiling against the deubiquitinases"

### Supplementary figures

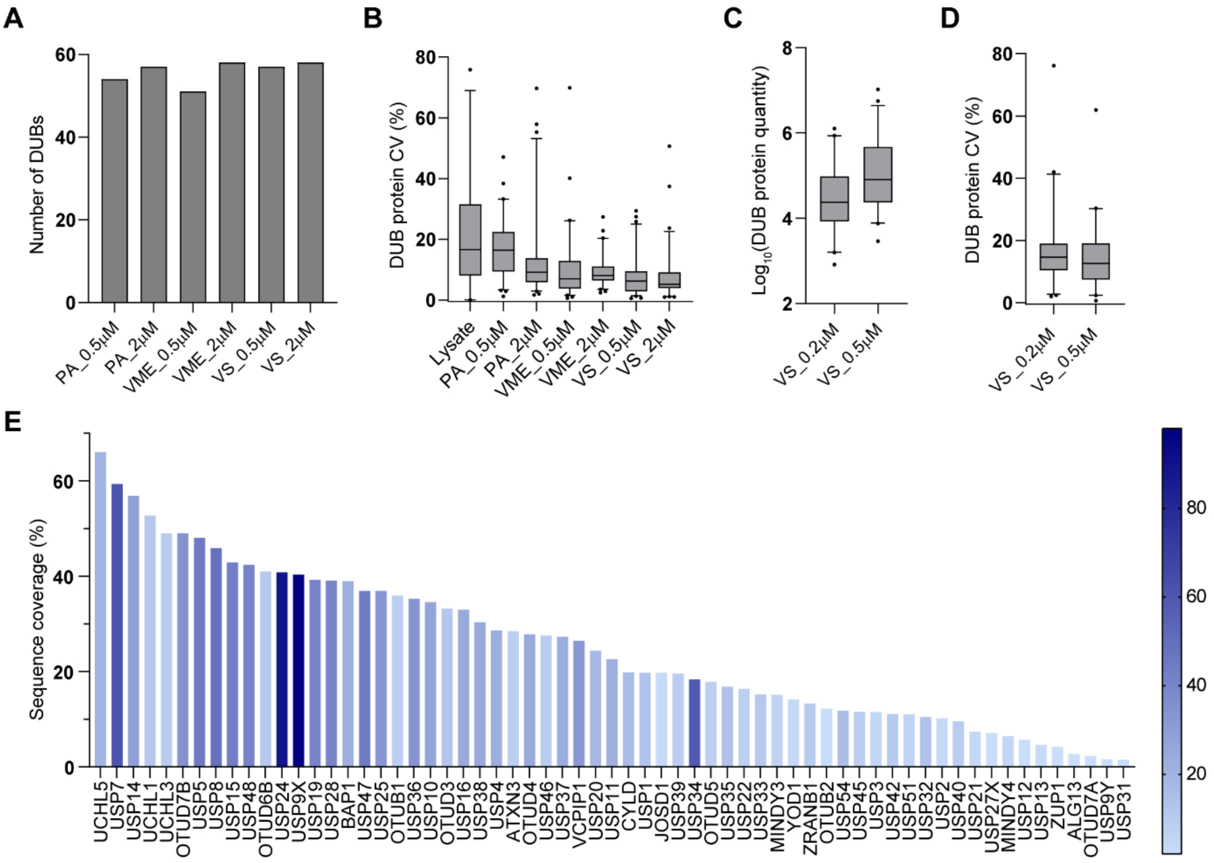

**SI figure 1.**

A) Number of significantly enriched DUBs when activity-based DUB probe treatment is compared to untreated lysate (avg log2 ratio ≤ -1, q-value ≤ 0.05).
B) Coefficient of variation (CV) of DUB protein quantities identified in at least two replicates (data from two biological replicates run in technical duplicates). Box plot shows the median central line and extends from 25$^{th}$ to 75$^{th}$ percentiles. Whiskers represent CVs within 5$^{th}$ to 95$^{th}$ percentiles.
C) DUB protein quantities to compare biotin-Ahx-Ub-VS treated samples at 0.5 and 0.2 µM. Box plot shows the median central line and extends from 25$^{th}$ to 75$^{th}$ percentiles. Whiskers represent protein quantity within 5$^{th}$ to 95$^{th}$ percentiles.
D) Coefficient of variation (CV) of DUB protein quantities identified in at least two replicates (data from two biological replicates run in technical duplicates). Box plot shows the median central line and extends from 25$^{th}$ to 75$^{th}$ percentiles. Whiskers represent CVs within 5$^{th}$ to 95$^{th}$ percentiles.
E) Bar chart showing DUB sequence coverage and number of identified peptides per quantified DUB for biotin-AhX-Ub-VS (0.5 µM) treated sample.

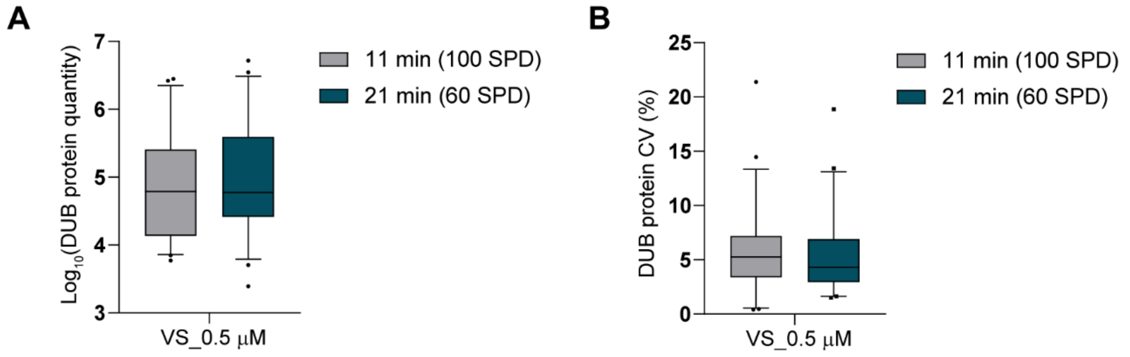

**SI figure 2. Comparison of 21 min and 11 min run times.**

A) DUB protein quantities for biotin-Ahx-Ub-VS (0.5 µM) treated samples to compare 11 min and 21 min run times. Box plot shows the median central line and extends from 25$^{th}$ to 75$^{th}$ percentiles. Whiskers represent protein quantity within 5$^{th}$ to 95$^{th}$ percentiles.

B) Coefficient of variation (CV) of DUB protein quantities identified in at least two replicates (data from three technical replicates). Box plot shows the median central line and extends from 25$^{th}$ to 75$^{th}$ percentiles. Whiskers represent CVs within 5$^{th}$ to 95$^{th}$ percentiles.

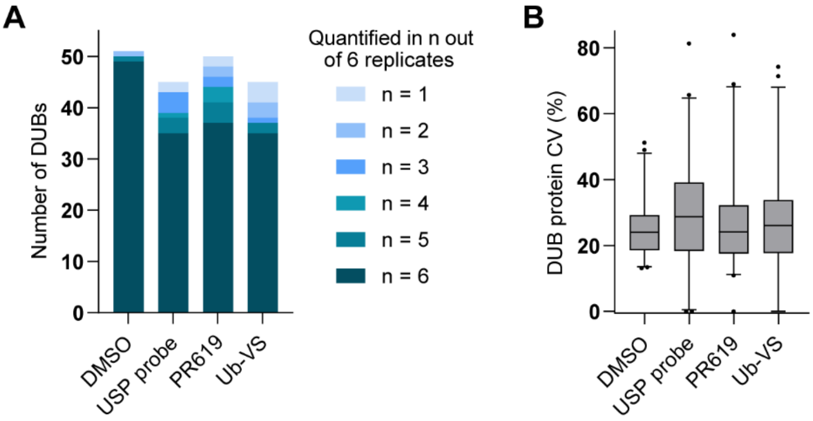

**SI figure 3. Reproducibility.**

A) Number of quantified DUBs in *n* out of six biological replicates.
B) Coefficient of variation (CV) of DUB protein quantities identified in at least two replicates (data from six biological replicates). Box plot shows the median central line and extends from 25$^{th}$ to 75$^{th}$ percentiles. Whiskers represent CVs within 5$^{th}$ to 95$^{th}$ percentiles.

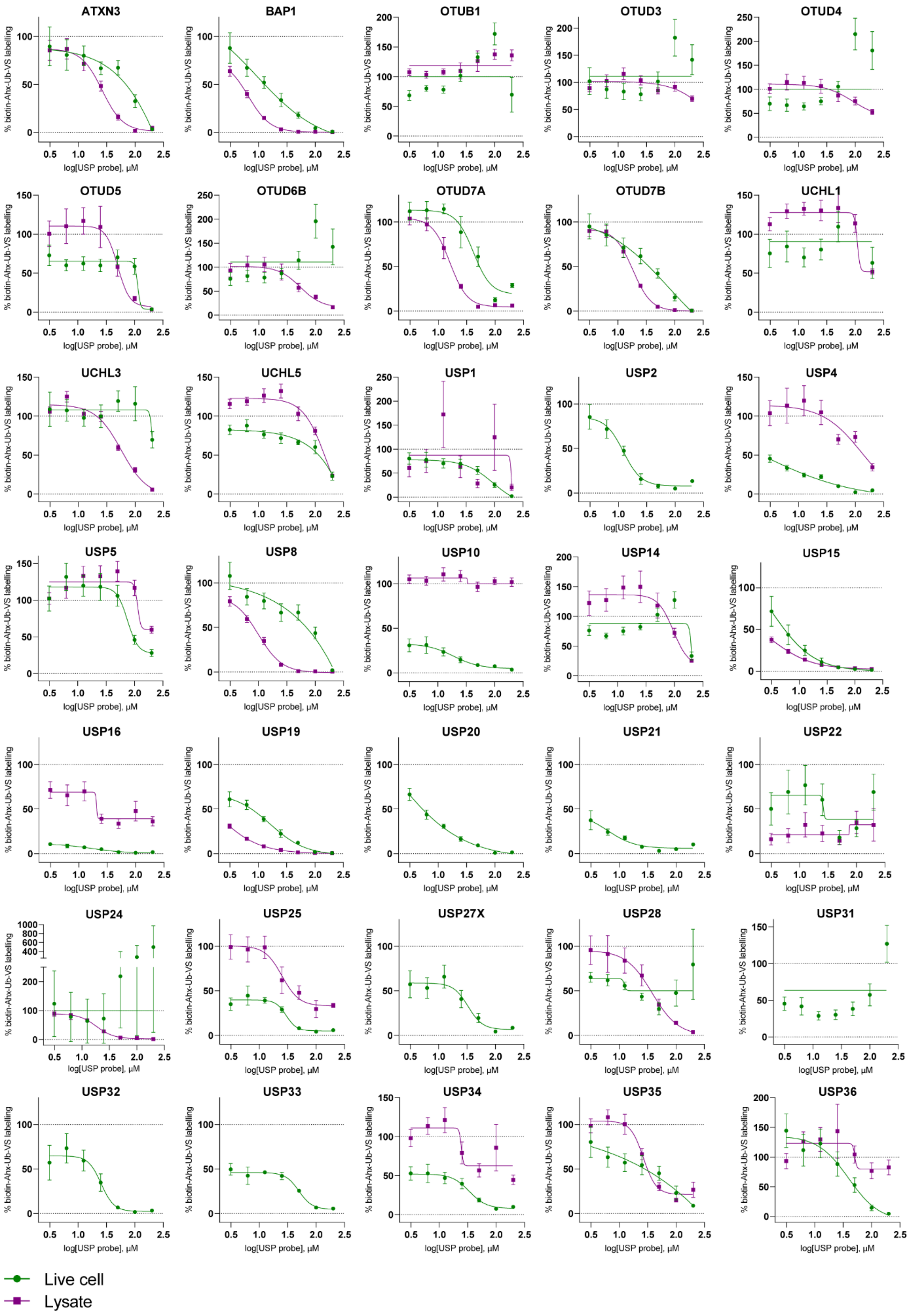
ATXN3
BAP1
OTUB1
OTUD3
OTUD4
OTUD5
OTUD6B
OTUD7A
OTUD7B
UCHL1
UCHL3
UCHL5
USP1
USP2
USP4
USP5
USP8
USP10
USP14
USP15
USP16
USP19
USP20
USP21
USP22
USP24
USP25
USP27X
USP28
USP31
USP32
USP33
USP34
USP35
USP36
% biotin-Ahx-Ub-VS labelling
log[USP probe], µM
Live cell
Lysate

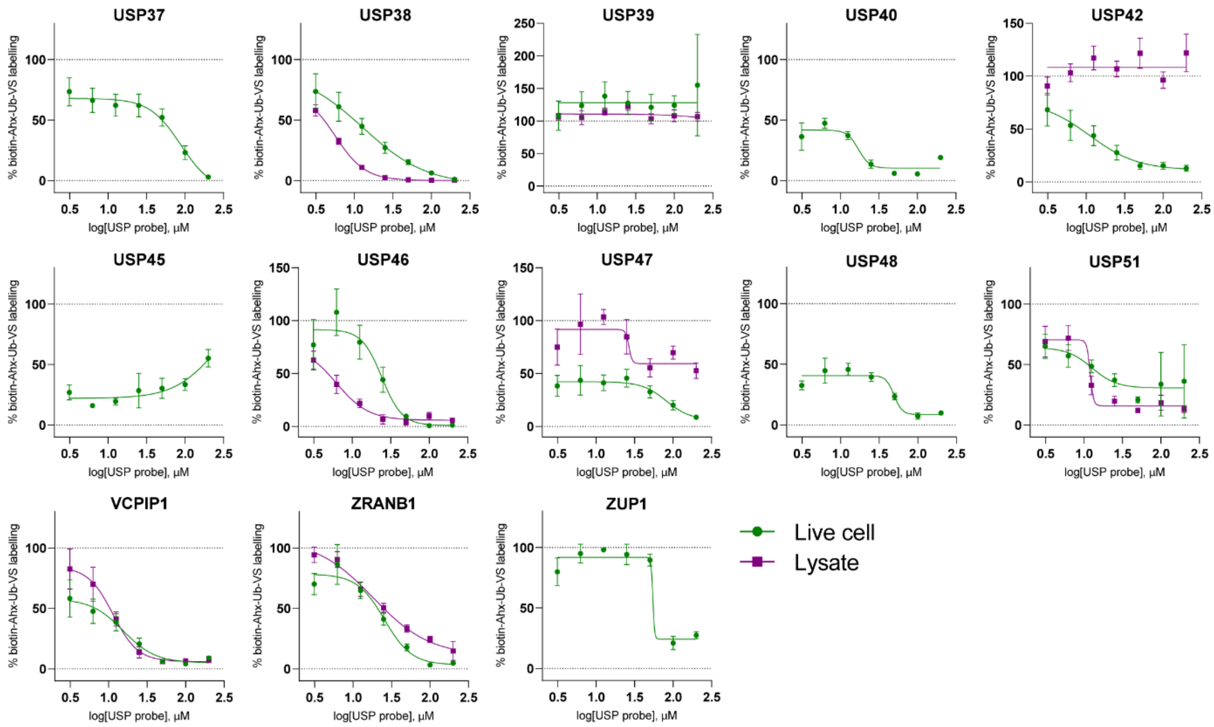

**SI figure 4. Rest of the live cell vs lysate dose response plots.** Data is presented as mean ± SEM, *n* = 3. The curves were fitted with GraphPad Prism 9 using four parameter nonlinear regression.

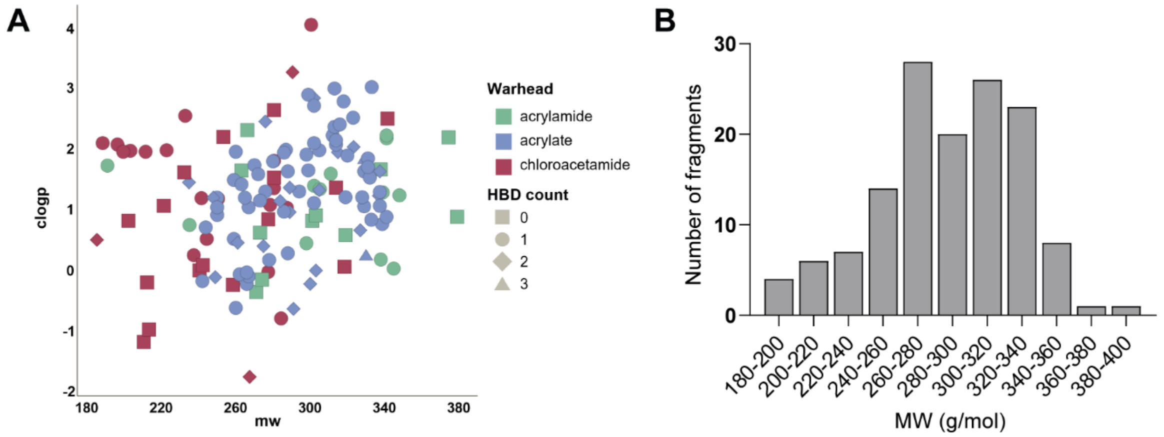

**SI figure 5. Fragment library**

A) Scatter plot of reactive fragment library showing distribution by mw and clogp, coloured by warhead, shaped by HBD count

B) Fragment library molecular weight distribution.

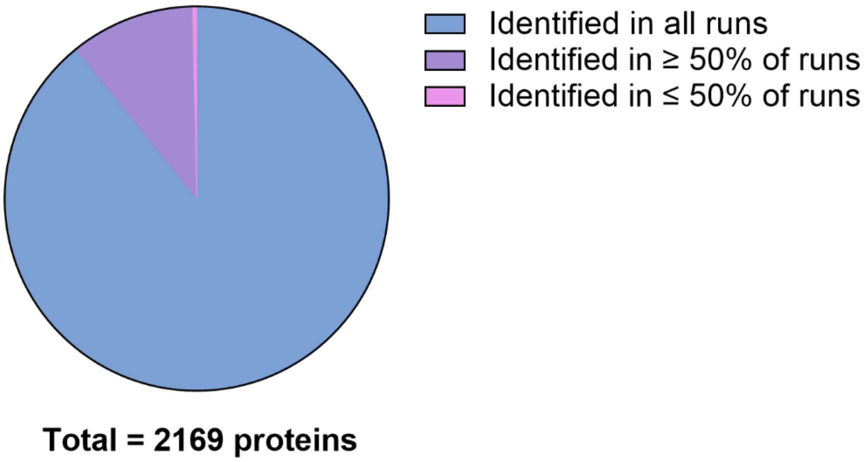

**SI figure 6. Instrument performance.** A pie chart showing the number of proteins identified in HEK293T cell lysate using 21 min predefined Evosep gradient (data from 24 technical replicates).

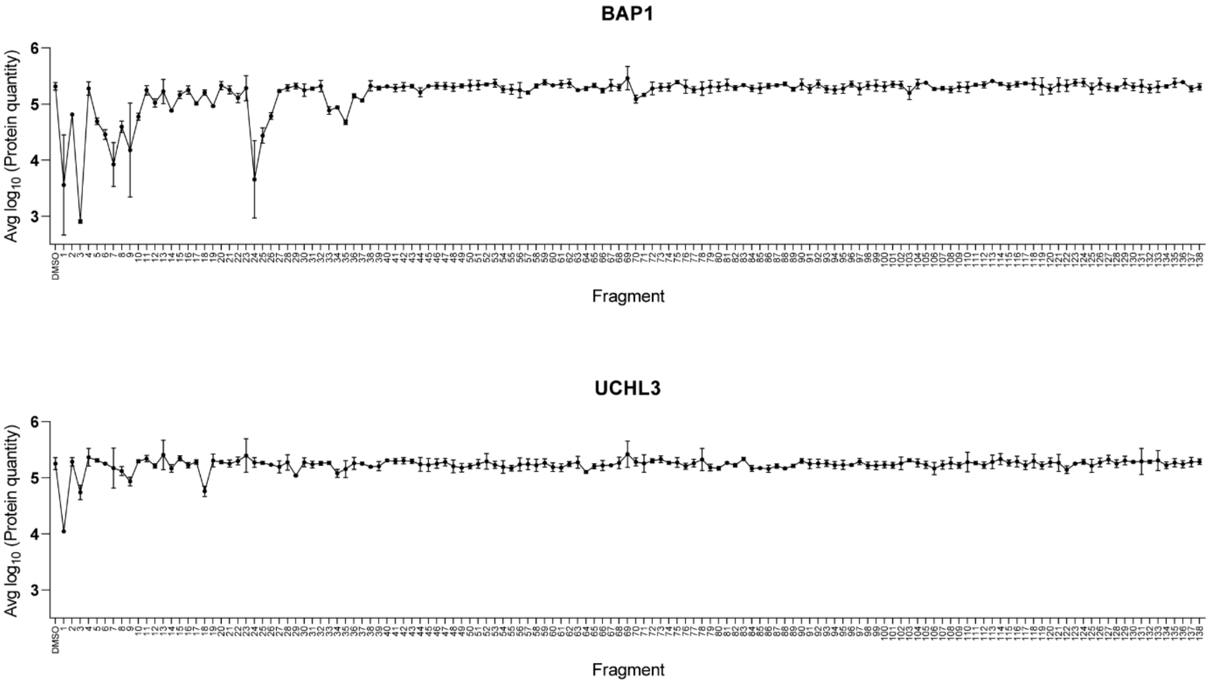

**SI figure 7. BAP1 and UCHL3 protein quantity data for DMSO and each fragment (200 µM).** Data is presented as mean ± SD, *n* = 3.

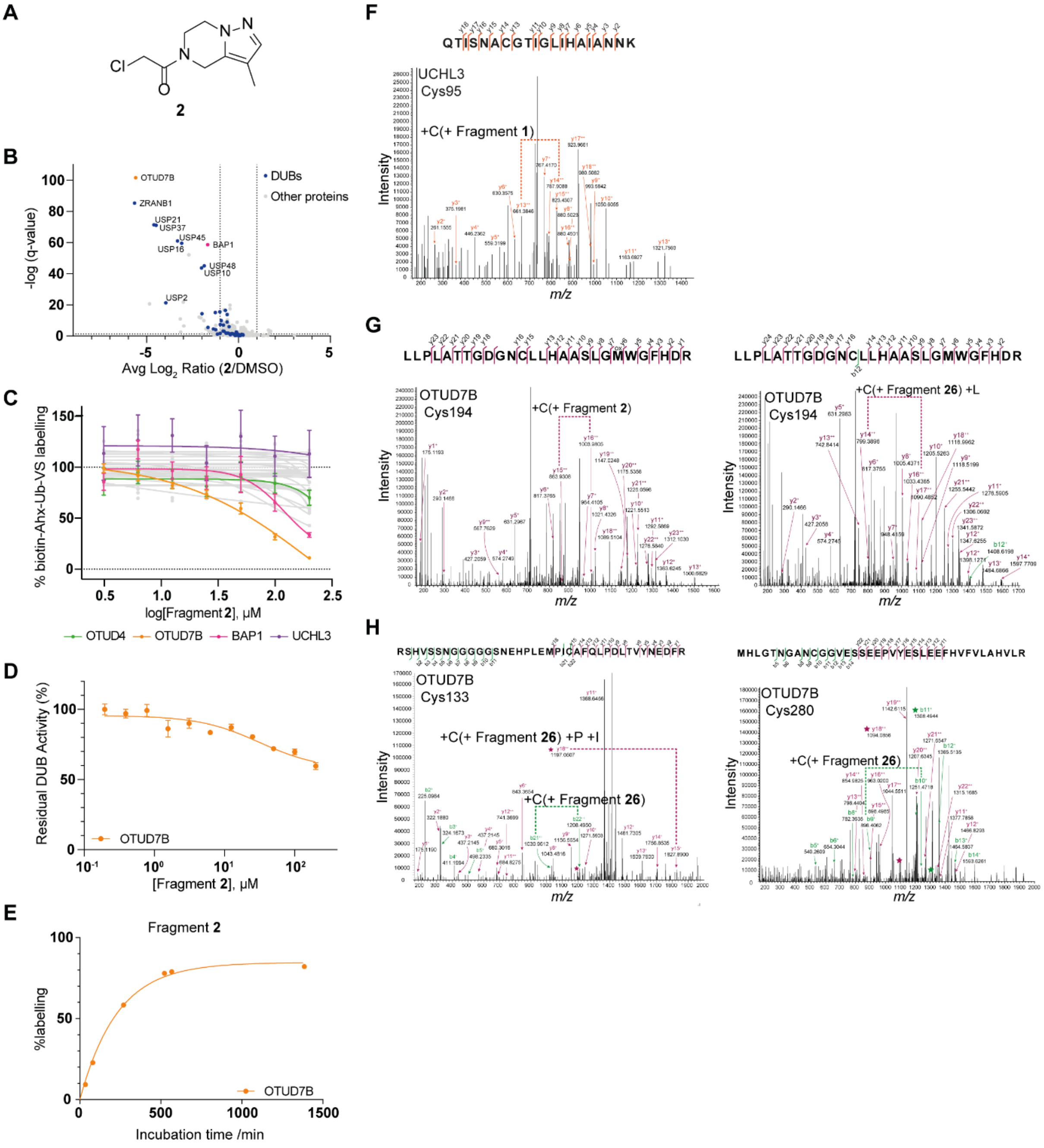

**SI figure 8. Hit fragment follow up**

A) Structure of hit fragment **2**.
B) Volcano plot showing significantly competed proteins (avg $\log_2$ ratio ≤ -1, q-value ≤ 0.05) when hit fragment **2** (200 µM) treated samples are compared to DMSO (biotin-Ahx-Ub-VS, 0.5 µM). DUBs are highlighted in blue.
C) Dose-response chemoproteomics data for hit fragment **2**. Data is presented as mean ± SEM, *n* = 3. The curves were fitted with GraphPad Prism 9 using four parameter nonlinear regression.
D) Enzymatic inhibition assay data for hit fragment **2**. Data is presented as mean ± sd, *n* = 3. The curve was fitted with GraphPad Prism 9 using three parameter nonlinear regression.
E) Intact-protein LCMS time-course %labelling data for fragment **2**. The curve was fitted with GraphPad Prism 9 using one phase association.
F) LC-MS/MS spectra of the peptide $_{89}$QTISNACGTIGLIHAIANNK$_{108}$ crosslinked to fragment **1** indicating UCHL3 Cys95 as the fragment binding site.

G) LC-MS/MS spectra of the peptide $_{183}$LLPLATTGDGNCLLHAASLGMWGFHDR$_{209}$ crosslinked to fragments **2** and **26** indicating OTUD7B Cys194 as the fragment binding site.

H) LC-MS/MS spectra of the peptides $_{111}$RSHVSSNGGGGGSNEHPLEMPICAFQLPDLTVYNEDFR$_{148}$ and $_{271}$MHLGTNGANCGGVESSEEPVYESLEEFHVFVLAHVLR$_{307}$ crosslinked to fragment **26** indicating OTUD7B Cys133 and Cys280 as minor fragment binding sites.

| DUB | USP probe lysate $IC_{50}$ (µM) | PR619 lysate $IC_{50}$ (µM) |
|---|---|---|
| **USP19** | 1.63 | 38.69 |
| **USP15** | 3.255 | 44 |
| **USP38** | 5.618 | ND |
| **USP46** | 6.215 | 4.31 |
| **BAP1** | 6.585 | 50.65 |
| **USP8** | 9.878 | 112.5 |
| **VCPIP1** | 11.37 | 5.789 |
| **USP51** | 12.01 | ND |
| **USP11** | 13.02 | 78 |
| **USP9X** | 13.48 | 6.343 |
| **OTUD7A** | 15.95 | 23.12 |
| **ZRANB1** | 18.08 | 58.75 |
| **USP24** | 18.25 | ND |
| **OTUD7B** | 18.32 | 19.68 |
| **USP34** | 24.34 | ND |
| **USP25** | 25.06 | 50.39 |
| **ATXN3** | 26.27 | 18.31 |
| **USP35** | 26.52 | 40.91 |
| **USP47** | 26.67 | ND |
| **USP7** | 31.97 | 48.42 |
| **USP28** | 38.05 | 39.25 |
| **OTUD5** | 50.47 | 2.225 |
| **USP36** | 50.59 | ND |
| **OTUD6B** | 53.57 | 14.61 |
| **UCHL3** | 57.17 | 32.58 |
| **OTUD4** | 90.48 | 41.41 |
| **USP14** | 95.29 | ND |
| **UCHL1** | 108.6 | 27.12 |
| **USP5** | 113 | 40.15 |
| **USP4** | 130 | 24.66 |
| **UCHL5** | 155.6 | 8.742 |
| **OTUD3** | 431.5 | 58.71 |
| **OTUB1** | ND | ND |
| **USP1** | ND | ND |
| **USP10** | ND | ND |
| **USP16** | ND | ND |
| **USP22** | ND | ND |
| **USP39** | ND | ND |
| **USP42** | ND | ND |
| **YOD1** | ND | 42.67 |

**SI Table 1:** $IC_{50}$ (µM) values for USP probe and PR619 in dose response lysate treatment for identified DUB proteins. $IC_{50}$ values were extracted from dose-response curves that were fitted with GraphPad Prism 9 using four parameter nonlinear regression. ND = Not determined

### Methods

#### KEY RESOURCES TABLE

| REAGENT or RESOURCE | SOURCE | IDENTIFIER |
|---|---|---|
| Bacterial and virus strains | | |
| NEB 5-alpha Competent E.coli cell line | New England Biolabs | Cat# C2987H |
| Competent BL21 GOLD (DE3) cell line | Agilent Technologies | Cat# 230132 |
| Competent BL21-CodonPlus (DE3) cell line | Agilent Technologies | Cat# 230280 |
| Chemicals, peptides, and recombinant proteins | | |
| Dulbecco's Modified Eagle Medium (DMEM) | Gibco | Cat#41966-029 |
| Fetal bovine serum (FBS) | Gibco | Cat#10270-106 |
| L-Glutamine–Penicillin–Streptomycin solution | Sigma Aldrich | Cat#G1146 |
| Biotin-Ahx-Ub-VS | UbiQ | Cat#UbiQ-188 |
| Biotin-Ahx-Ub-VME | UbiQ | Cat#UbiQ-054 |
| Biotin-Ahx-Ub-PA | UbiQ | Cat#UbiQ-076 |
| Ub-VS | UbiQ | Cat#UbiQ-108 |
| Pierce™ High Capacity NeutrAvidin™ Agarose | Thermo Fisher Scientific | Cat#29204 |
| Ub-Rho110Gly | UbiQ | Cat#UbiQ-002 |
| HisPur™ NiNTA resin | Thermo Fisher Scientific | Cat#88222 |
| Glutathione Sepharose 4B | Sigma Aldrich | Cat#GE17-0756-01 |
| Compound 1 | Enamine | Cat#Z295410654 |
| Compound 2 | Enamine | N/A |
| Compound 3 | Enamine | Cat#Z57042000 |
| Compound 26 | Enamine | Cat#Z3766262724 |
| USP Probe | Ward et al., 2016 | N/A |
| PR-619 | Sigma Aldrich | Cat# SML0430 |
| Critical commercial assays | | |
| Pierce™ BCA Protein Assay Kit | Thermo Scientific | Cat#23227 |
| Deposited data | | |
| Raw and analyzed proteomics data | This paper | PXD037114 and PXD037152 |
| Experimental models: Cell lines | | |
| HEK293T cell line | ATCC | Cat# CRL-3216 |
| Recombinant DNA | | |
| pGEX6P2 UCH-L3 plasmid | MRC PPU Reagents and Services | DU Number 21025 |
| UCHL1 cDNA (BC005117) | imaGenes (www.imagenes-bio.de) | Cat# IRAUp969F0918D |
| pOPINK-Cezanne (OTU, aa 53-446) (OTUD7B) plasmid | Mevissen et al., 2013 | Addgene plasmid #61581 |
| Software and algorithms | | |
| Spectronaut 15.4.210913.50606 | Biognosys | https://biognosys.com/software/spectronaut/ |
| GraphPad Prism 9.4.0 | GraphPad | https://www.graphpad.com/ |

| | | |
|---|---|---|
| Mascot 2.8.0 | Matrix Science | https://www.matrixscience.com/ |
| MaxQuant 1.6.12.0 | Cox, J., and Mann, M. (2008). MaxQuant enables high peptide identification rates, individualized p.p.b.-range mass accuracies and proteomewide protein quantification. Nat. Biotechnol. 26, 1367–1372. | https://maxquant.net/maxquant/ |
| BioRender | BioRender | www.biorender.com |
| Other | | |
| Combinatorial Microlute™ plate | Porvair | Cat#24002 |

**RESOURCE AVAILABILITY**

**Lead contact**

Further information and requests for resources and reagents should be directed to and will be fulfilled by the lead contacts, Jacob Bush and Katrin Rittinger.

**Materials availability**

Recombinant proteins were either expressed using commercially available plasmids or DNA. Please see key resources table below for further information. Compounds are commercially available from Enamine or Sigma Aldrich, as stated in the key resources table.

**Data and code availability**

The raw mass spectrometry proteomics files and database search results have been deposited at the ProteomeXchange Consortium (http://proteomecentral.proteomexchange.org) via the PRIDE partner repository (Perez-Riverol et al. The PRIDE database and related tools and resources in 2019: improving support for quantification data. Nucleic Acids Res. 2019 Jan 8;47(D1):D442-D450.) with data set identifier PXD037114 and PXD037152.

**EXPERIMENTAL MODEL AND SUBJECT DETAILS**

**Cell lines**

HEK293T cell line (female human origin, ATCC, Cat# CRL-3216) was used in this study. The cells were maintained at 37 °C with 5% $CO_2$ in DMEM media supplemented with 10% fetal bovine serum and 1% L-Glutamine–Penicillin–Streptomycin solution (200 mM L-glutamine, 10 000 U/mL penicillin and 10 mg/mL streptomycin). Cells were authenticated by STR profiling by Francis Crick Institute Cell Services STP. All cells were negative from mycoplasma contamination tested by fluorescence staining, agar culture, and PCR testing by Francis Crick Institute Cell Services STP.

#### METHOD DETAILS

##### Acetylation of NeutrAvidin agarose beads

20 mL of high capacity NeutrAvidin agarose beads were centrifuged (2000 rcf, 2 min, RT) and supernatant was removed. The beads were washed three times with 15 mL of PBS. 8.8 mL of PBS and 482 µL of freshly made NHS-acetate (100 mg, 20 mM final concentration in anhydrous DMSO) were added. The beads were incubated for 30 min at room temperature on rocker. Further 482 µL of freshly made NHS-acetate (100 mg, in anhydrous DMSO) was added and the incubation step was repeated. The reaction was quenched by adding 4 mL of 1 M Tris pH 7.5. The beads were washed twice with 15 mL of PBS and twice with 15 mL of 20% EtOH. 10 mL of 20% EtOH was added and the beads were stored at 4 °C until needed.

##### Lysate preparation and treatment

HEK293T cell pellets (pre-washed twice with PBS) were lysed in lysis buffer containing 50 mM TRIS pH 7.5, 150 mM NaCl, 0.5% CHAPS hydrate, 0.1% IGEPAL CA-360, 5 mM MgCl2, 10% Glycerol and protease inhibitor cocktail (1:100) by using Branson probe sonicator (5 x 1 sec, 10% amplitude). The lysate was clarified by centrifugation (5000 rpm, 20 min, 4 °C). The protein concentration was determined using a BCA assay and lysate was diluted to 2 mg/mL.

244 µL aliquots of lysate were incubated for 3 h with 5 µL of fragments/inhibitors or DMSO at room temperature followed by 1 h incubation with biotin-Ahx-Ub-Vs (0.5 µM final concentration). Lysates were diluted to 0.6 mg/mL with 0.2% SDS in PBS. All treatments were done at least in two biological replicates and the number of replicates and treatment concentrations for each experiment are stated in figure legends or results section.

##### Live cell treatment

HEK293T cells were treated for 3 h in triplicate with USP probe (3.125-200 µM) or DMSO vehicle at 37 °C in serum free media. Media was removed and cells were washed with PBS. Cells were lysed in lysis buffer containing 50 mM TRIS pH 7.5, 150 mM NaCl, 0.5% CHAPS hydrate, 0.1% IGEPAL CA-360, 5 mM $MgCl_2$, 10% Glycerol and protease inhibitor cocktail (1:100) and benzonase (1:1000). The lysates were clarified by centrifugation (16 000 rcf, 10 min, 4 °C). Protein concentration of each lysate was determined using a BCA assay and diluted to 2 mg/mL. The lysates were incubated for 1 h with biotin-Ahx-Ub-Vs (0.5 µM final concentration) at room temperature. Lysates were diluted to 0.6 mg/mL with 0.2% SDS in PBS.

##### Enrichment and digestion

Samples were incubated with 60 µL of acetylated NeutrAvidin agarose beads (pre-washed three times with 1:3 lysis buffer:0.2% SDS in PBS) on combinatorial microlute filtration plate for 2 h. Supernatants were removed by centrifugation (700 rcf, 1 min) and the beads were washed three times with lysis buffer, 4 M urea in 50 mM HEPES pH 8.5 and 50 mM HEPES pH 8.5.

NeutrAvidin beads were incubated for 30 min with 5 mM tris(2-carboxyethyl)phosphine (TCEP) and 15 mM 2-Chloroacetamide (CAA) in 50 mM HEPES pH 8.5. Supernatants were removed by centrifugation (700 rcf, 1 min) and the beads were washed with 50 mM HEPES pH 8.5.

The proteins were digested on-bead overnight at 37 °C with 60 µL of LysC (0.004 µg/µL) in 50 mM HEPES pH 8.5. The supernatants were collected by centrifugation (700 rcf, 1 min) and 40 µL of trypsin (0.006 µg/µL) in 50 mM HEPES pH 8.5 was added to each sample. The samples were digested for 4 h at 37 °C and acidified with formic acid.

**Preparation of digested HEK293T cell lysate**

HEK293T lysate (2 mg/ml) was incubated for 30 min with 5 mM tris(2-carboxyethyl)phosphine (TCEP) and 15 mM 2-Chloroacetamide (CAA) in 50 mM HEPES pH 8.5. Proteins were precipitated using ice-cold acetone and the resulting pellets were washed twice with ice-cold 80% acetone. The air-dried pellets were dissolved in 1 M guanidinium chloride in 50 mM HEPES pH 8.5 by vortexing and sonicating. The proteins were digested overnight at 37 °C with LysC in 50 mM HEPES pH 8.5 followed by 4 h digestion with trypsin at 37 °C. 1 µg of digested lysate was loaded to Evotips as described below.

**LC-MS/MS analysis**

The samples were loaded with iRT standard onto Evotips (as prepared according to manufacturer's instructions) followed by loading onto the Evosep One LC system in front of the Orbitrap Fusion Lumos Tribrid mass spectrometer (Thermo Fisher Scientific). Data for all samples was acquired in Data Independent Acquisition mode (DIA) using the pre-set gradients on Evosep One.

For the 44 minute predefined method Evosep One was fitted with a 15 cm column (EV1113). DIA Lumos settings were as follows: Transfer capillary set to 300 °C and 2.2kV applied to the nanospray needle (Evosep). MS1 data acquired in the Orbitrap with a resolution of 120k, max injection time of 20 ms, AGC target of 1e6, in positive ion mode, in profile mode, over the mass range 393-907 *m/z*. DIA segments over this mass range (20 *m/z* wide/1 Da overlap/27 in total) were acquired in the Orbitrap following fragmentation in the HCD cell (32%), with 30k resolution over the mass range 200-2000 *m/z* and with a max injection time of 54ms and AGC target of 1e6.

For 21 and 11 minute methods Evosep One was fitted with an 8 cm column (EV1064).

21 minute method used the same settings with the following changes: MS1 data was acquired over the mass range 392-811 *m/z*. DIA segments over this mass range were set as 17.5 *m/z* wide/1 Da overlap/25 in total.

11 min method used the same settings with the following changes: MS1 data was acquired over the mass range 392-811 *m/z*. DIA segments over this mass range were set as 21 *m/z* wide/1 Da overlap/19 in total.

The raw data was searched and analysed using Spectronaut (v. 15.4.210913.50606) against human uniprot (August 2019), contaminants and avidin fasta files using directDIA method. QUANT 2.0 was selected as Protein LFQ Method. The data was normalised using global median normalisation strategy with automatic row selection. Run wise imputation with Q-value percentile = 10% was selected. Other search settings were used as default (BGS factory settings). Two sample t-test was used to assess average $\log_2$ ratios (fragment/DMSO). Before plotting all data was filtered for unique peptides ≥ 2. Significantly competed DUBs were defined based on average $\log_2$ ratio ≤ -1 and q-value ≤ 0.05.

The graphs were plotted and curves were fitted using GraphPad Prism (v. 9).

**Recombinant Protein Expression and Purification**

Recombinant UCHL3 was expressed and purified from BL21 E.coli cells. OTUD7B(53-446) was expressed and purified from CodonPlus BL21 E.coli cells. The proteins were purified via GST or His-tag enrichment, before tag-cleavage with 3C protease. The DUBs were further purified through gel filtration chromatography and stored in 50 mM HEPES pH 7.5, 150 mM NaCl, 0.5 mM TCEP and 5% glycerol.

**DUB Inhibition Assay**

Prior to inhibition assays, optimal conditions for each DUB were found by performing plate-based fluorescence time courses with a matrix of DUB and substrate dilutions. All assays were performed in 50 mM HEPES pH 7.5, 100 mM NaCl, 1 mM EDTA, 0.05% Tween20, 10 mM DTT. The reactions were initiated with the addition of Ub-Rho110Gly and monitored by measuring the rhodamine fluorescence every 45 s at room temperature in a Clariostar Plus plate reader.

DUBs (2.5 µM) were pre-treated with chosen fragments (0-200 µM) for 3 h at room temperature. Following dilution of the DUBs, reactions were initiated with the addition of Ub-Rho110Gly and monitored as above. Final reaction concentrations were as follows: 62.5 nM Ub-Rho110Gly with either 12 pM UCHL3 or 12.5 nM OTUD7B. Initial reaction rates were calculated by taking away background fluorescence of Ub-Rho110 hydrolysis and plotting fluorescence against time. Reaction rates were normalised against samples treated with the lowest concentration of fragment and plotted as percentages of residual DUB activity against fragment concentration using GraphPad Prism (v. 9). The curves were fitted using three parameter nonlinear regression. Each experiment was set up as technical triplicates.

**Intact Protein LC-MS**

0.25 µM stock solutions of recombinant OTUD7B and UCHL3 were prepared in assay buffer (25 mM HEPES pH 7.5, 50 mM NaCl). 80 µL of protein stocks for each time point were pipetted into 384-well plate and 0.4 µl of fragment (50 µM final concentration) or DMSO was added. The plate was incubated for 24 h and reaction was measured by LC-MS over six time points (29, 74, 264, 516, 561 and 1380 min) as described below.

Intact protein masses were recorded by LC-MS using an Agilent G6230B time-of-flight (ToF) Accurate Mass Series mass spectrometer, interfaced with an Agilent 1290 infinity II series column oven (G7116B) and an Agilent 1290 infinity II series liquid chromatography high speed binary pump (G7120A). The protein sample was injected using an Agilent 1290 infinity II series multisampler with dual needles (Model No. G7167B) with a 10 µL injection volume and maintained at a temperature of 4 °C. Chromatography was carried out on an Agilent Bio-HPLC PLRP-S (1000 Å, 5 µm × 50 mm × 1.0 mm, PL1312-1502) reverse phase HPLC column at 70 °C. The sample was eluted at 0.5 mL/min using a gradient system from Solvent A (water, 0.2% (*v/v*) formic acid) to Solvent B (acetonitrile, 0.2% (*v/v*) formic acid) according to the conditions described in Table S13. The eluent was injected directly into an Agilent ToF mass spectrometer (Model No. G6230B) using a dual AJS ESI source and scanning between 600–3200 Da with a scan rate of 1.20 s in positive mode. The following MS parameters were used: 4000 V capillary voltage limit, 350 °C desolvation temperature, 10 L/min drying gas flow.

SI Table 1 - Intact-protein LCMS solvent gradient.

| Time (min) | Flow rate / mL/min | Solvent A % | Solvent B % |
|---|---|---|---|
| 0.60 | 0.5 | 80 | 20 |
| 0.61 | 0.5 | 50 | 50 |
| 1.00 | 0.5 | 0 | 100 |
| 1.20 | 0.5 | 0 | 100 |
| 1.21 | 0.5 | 80 | 20 |

Data acquisition was carried out in 2 GHz Extended Dynamic range mode. Spectra were processed using Mass Hunter Qualitative Analysis™ B06.00 (Agilent) software with the Maximum Entropy method employed. The total ion chromatograms (TIC) were extracted (region containing protein) and the summed scans were deconvoluted (using a maximum entropy algorithm) over a *m/z* range with an expected mass range dependent on the protein.

SI Table 2. The deconvolution conditions for recombinant proteins studied in this work.

| **Protein** | **Expected mass range** | ***m/z* range** |
|---|---|---|
| OTUD7B | 40000-50000 | 800–2000 |
| UCHL3 | 10000-50000 | 800–2000 |

The peak height for unmodified and modified protein were recorded from Bioconfirm and used to calculate percentage of labelling for each time point using **Error! Reference source not found.**.

% = ((intensity of modified protein)/(intensity of protein only+(intensity of modified protein)))*100

Equation 1. Relating the percentage crosslinking to the intensities of protein and modified protein, as measured by intact protein LCMS.

Percentage of labelling was plotted against incubation time for all compounds and proteins using GraphPad Prism (v. 5) and one phase association model with y0 constrained to = 0.

**Site ID experiment**

1 µM solution of recombinant OTUD7B and UCHL3 were prepared in assay buffer (25 mM HEPES pH 7.5, 50 mM NaCl). Protein solutions were incubated with fragments (100 µM final concentration) at room temperature before being flash frozen after 8 h.

Reaction mixtures were thawed and incubated with iodoacetamide (1 mM final concentration) for 30 min in the dark. 20 µL of freshly prepared S-Trap lysis buffer (5% SDS, 50 mM TEAB, pH 8.5) was added and samples were vortexed briefly to ensure mixing. A further addition of iodoacetamide along with TCEP was performed to a final concentration of 10 mM of each and samples were incubated for 30 min in the dark. Samples were acidified by addition of 2.5 µL phosphoric acid (25%) before addition of 165 µL of S-Trap binding buffer (100 mM TEAB, 90% MeOH, pH 7.55). Samples were vortexed to ensure mixing and precipitation of proteins. 120 µL of samples were loaded onto individual S-Trap micro columns followed by centrifugation (4000 rcf, 30 s). Loading of 120 µL volumes was repeated until all the samples were loaded onto the columns. Trapped protein samples were washed by addition of 150 µL of S-Trap binding buffer followed by centrifugation (4000 rcf, 30 s). This step was repeated for a total of four times. For OTUD7B and UCHL3 samples, 20 µL of trypsin (0.05 µg/µL) was added to each column and samples were incubated at 47 °C for 2 h. For BAP1, 20 µL of AspN (0.05 µg/µL) was added and samples were incubated at 47 °C for 1 h. AspN solution was centrifuged (4000 rcf, 30 s) through the column at and discarded before addition of 20 µL of trypsin (0.05 µg/µL) to columns. Samples were incubated at 47 °C for a further 2 h. Peptides were eluted from S-Trap columns by subsequent addition and centrifugation (4000 rcf, 30 sec) of 40 µL of TEAB 50 mM, 40 µL of 0.1% formic acid and 40 µL of 0.1% formic acid, 50% acetonitrile. Eluted peptides were flash frozen on dry ice before vacuum drying.

Dried peptides were resuspended in 0.1% formic acid to a final concentration of 50 ng/µL and 6 µL was injected onto an Orbitrap Fusion Lumos Tribrid mass spectrometer.

Peptide samples were separated on an Ultimate 3000 RSLC system (Thermo Fisher Scientific) with a C18 PepMap, serving as a trapping column (2 cm × 100 µm ID, PepMap C18, 5 µm particles, 100 A pore size) followed by a 50 cm EASY-Spray column (50 cm × 75 µm ID, PepMap C18, 2-µm particles, 100 A pore size) (Thermo Fisher Scientific). Buffer A contained 0.1% FA and Buffer B 80% MeCN, 0.1% FA. Peptides were separated first by holding 2% Buffer B for 6 min before a step to 8% B in 1 min Peptides were then separated with a linear gradient of 8–45% (Buffer B) over 40 min followed by a step from 45 to 95% Buffer B 2 min at 250 nL/min and held at 90% for 5.5 min. The gradient was then decreased to 2% Buffer B in 0.5 min at 250 nL/min for 15 min.

Mass spectrometric analysis was performed on an Orbitrap Fusion Lumos Tribrid mass spectrometer (Thermo Fisher Scientific) operated in "TopSpeed" data dependant mode in positive ion mode with a 3 sec cycle time. Full scan spectra were acquired in a range from 375 to 1500 m/z, at a resolution of 120000 (at 200 *m/z*), with an automatic gain control (AGC) target of standard and a maximum injection time set to auto. Charge states included for analysis were 2-5$^+$ ions. For MS/MS analysis, the Orbitrap resolution was set to 50000 (at 200 *m/z*). A minimum AGC target was set to standard and the most intense precursor ions were isolated with a quadrupole mass filter width of 1.2. Precursors were subjected to higher-energy collisional dissociation (HCD) fragmentation that was performed using a stepped-step collision energy of 25, 29 and 32%.

Uninterpreted tandem MS spectra were searched for peptide matches against the human sequences of OTUD7B and UCH3L along with common contaminants. Raw files were searched with both Mascot software (Matrix Science, Version 2.8.0) as well as MaxQuant (Andromeda, Version 1.6.12.0) [Cox J, et al, *Nature Biotechnology,* **26**, 2008]. A 4.5-5 ppm mass tolerance for peptide precursors and 20-25 ppm mass tolerance for fragment ions were selected.

OTUD7B and UCH3L raw files were searched using trypsin as the enzyme. Up to 2 missed cleavages were allowed and the variable modifications carboamidomethylation (C) and oxidation (M) were included. Compound specific modifications were added to the relevant databases and allowed on cysteine residues only and added to searches as variable modifications. Validation of binding was performed with manual interpretation of raw spectra.

**Chemistry**

*PR-619 was purchased from Sigma Aldrich. Compounds 1, 2, 3, and 26 were purchased from Enamine.*

*USP probe was synthesised from commercial reagents as desribed below.*

**N-benzyl-4-(2-chloroacetyl)-1-methyl-1H-pyrrole-2-carboxamide (USP probe)**

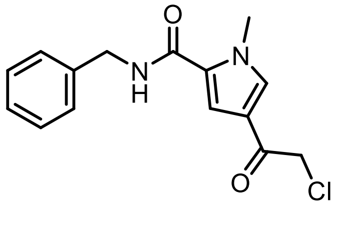

To a solution of 4-(2-chloroacetyl)-1-methyl-1H-pyrrole-2-carboxylic acid (200 mg, 0.96 mmol) in THF (8 mL) was added EDC (221 mg, 1.16 mmol), HOBt (154 mg, 0.96 mmol), followed by benzylamine (0.126 mL, 1.16 mmol). The reaction was stirred at room temperature for 3.5 hr. The reaction mixture was diluted in EtOAc (15 mL), washed with water (1 x 15 mL), organic layer separated and dried over anhydrous sodium sulfate, filtered and concentrated under reduced pressure to afford the crude product as a yellow gummy solid. Solid was purified by Biotage Isolera column chromatography (0-100% EtOAc in petroleum ether). Fractions containing compound were collected and concentrated under reduced pressure too afford desired product as an off-white solid (70 mg, 0.24 mmol, 24.9%). **LCMS** $t_r$ = 0.89 min, purity = 99% by UV, [M+H] = 291.0 Da. **$^1$H NMR** (400 MHz, DMSO-d6): δ 8.86 (t, J = 6.00 Hz, 1H), 7.87 (d, J = 1.60 Hz, 1H), 7.35-7.23 (m, 6H), 4.77 (s, 2H), 4.40 (d, J = 6.00 Hz, 2H), 3.90 (s, 3H).

#### QUANTIFICATION AND STATISTICAL ANALYSIS

Details of replicates and data analysis for specific experiments can be found in figure legends or the methods section.
